# Loss of adipocyte phospholipase gene *PLAAT3* causes lipodystrophy and insulin resistance due to inactivated arachidonic acid-mediated PPAR*γ* signaling

**DOI:** 10.1101/2021.04.15.439941

**Authors:** Nika Schuermans, Salima El Chehadeh, Dimitri Hemelsoet, Elke Bogaert, Elke Debackere, Pascale Hilbert, Nike Van Doninck, Marie-Caroline Taquet, Toon Rosseel, Griet De Clercq, Carole Van Haverbeke, Jean-Baptiste Chanson, Benoit Funalot, François-Jérôme Authier, Sabine Kaya, Wim Terryn, Steven Callens, Bernard Depypere, Jo Van Dorpe, Program for Undiagnosed Diseases (UD-PrOZA), Bruce Poppe, Christel Depienne, Bart Dermaut

**Affiliations:** Center for Medical Genetics, Ghent University Hospital, Ghent, 9000, Belgium; Department of Biomolecular Medicine, Faculty of Medicine and Health Sciences, Ghent University, Ghent, 9000, Belgium; Service de Génétique Médicale, Hôpital de Hautepierre, Strasbourg, 67000, France; Department of Neurology, Ghent University Hospital, Ghent, 9000, Belgium; Institute of Pathology and Genetics, Department of Molecular and Cell Biology, Charleroi, 6041, Belgium; General Hospital AZ Nikolaas, Department of Endocrinology and Diabetology, Sint-Niklaas, 9100, Belgium; Department of Internal Medicine and Nutrition, Hopitaux Universitaires Strasbourg, Strasbourg, 67000, France; Department of Pathology, Ghent University Hospital, Ghent, 9000, Belgium; Service de neurologie Hôpital de Hautepierre, Strasbourg, 67000, France; Department of Medical Genetics, Hôpital Henri Mondor, Université Paris-Est-Créteil, Créteil, 94010, France; Inserm UMR955, Créteil, 94010, France; Centre Expert de Pathologie Neuromusculaire, Hôpital Henri Mondor, Université Paris-Est-Créteil, Créteil, 94010, France; Institut für Humangenetik, Universitätsklinikum Essen, Essen, 45147, Germany; Department of Nephrology, Jan Yperman Hospital, Ieper, 8900, Belgium; Department of General Internal Medicine, Ghent University Hospital, Ghent, 9000, Belgium; Department of Plastic and Reconstructive Surgery, Ghent University Hospital, Ghent, 9000 Belgium

**Author notes:** These authors contributed equally to this work (Nika Schuermans, Salima El Chehadeh, Dimitri Hemelsoet). **Collaborators:** Program for Undiagnosed Diseases (UD-PrOZA): Steven Callens, Paul Coucke, Bart Dermaut, Dimitri Hemelsoet, Bruce Poppe, Nika Schuermans, Wim Terryn, Rudy Van Coster, Arnaud Vanlander, Patrick Verloo.

## Abstract

PLAAT3 is a phospholipid modifying enzyme predominantly expressed in white adipose tissue (WAT). It is a candidate drug target as Plaat3 deficiency in mice protects against picornavirus infection and diet-induced obesity. We identified four patients with homozygous loss-of-function mutations in *PLAAT3*, presenting with partial lipodystrophy, severe insulin resistance and dyslipidemia. PLAAT3-deficient WAT showed a failure to liberate arachidonic acid (AA) from membrane phospholipids resulting in an inactive gene network downstream of adipogenesis master regulator and anti-diabetic drug target PPARG. These findings establish PLAAT3 deficiency in humans as a novel type of partial lipodystrophy due to an AA- and PPARG-dependent defect in WAT differentiation and function.

## Background

Human PLAAT3, previously known as phospholipase A2, group XVI (PLA2G16), is part of a small family of acyltransferases and phospholipases related to lecithin:retinol acyl transferase (LRAT) which catalyze phospholipase A (PLA) and acyltransferase (AT) activities.^1^ In mice, Plaat3 is highly expressed in WAT where it mainly exhibits PLA2 activity which hydrolyzes fatty acids such as arachidonic acid (AA) linked to the *sn*-2 positions of membrane phospholipids.^1-3^ Plaat3 deficient mice are protected against picornavirus infection and are resistant to diet-induced obesity.^4, 5^ PLAAT3 has not yet been associated with a phenotype in humans, but it has been studied as a diagnostic or prognostic biomarker for different types of malignancies.^6-8^ Here, we undertook an integrated clinical, genetic and multi-omics approach in two consanguineous families to identify genetic PLAAT3 deficiency as a novel cause of partial lipodystrophy caused by an AA- and PPARG-mediated defect in WAT differentiation.

## Methods

### Patient clinical characteristics

All four patients were clinically evaluated by an endocrinologist and a neurologist. Blood samples were taken for dosage of fasting glucose, HbA1c, insulin, cholesterol, triglycerides and leptin. Plasma insulin measurements were done using the Roche ECLIA technology, leptin was measured through a radioimmunoassay. Lipoatrophy and regional fat accumulation were objectified using MRI or CT scan imaging. In one patient a dual X-ray absorptiometry was performed to evaluate body composition. In two patients an EMG and a brain MRI were done. In one patient and one control, biopsy specimens of upper arm subcutaneous WAT were obtained for histopathological examination. Quantification of the results was performed using the ImageJ software.^9^ Statistical analysis was done in SPSS Statistics 27. The study was approved by the Ethics Committee of the University Hospital of Ghent (EC: 2019/1430). Written informed consents for multi-omics analysis and publication of clinical pictures were obtained from all patients and control individuals.

### Genomics

Genetic studies in family 1 were initiated by the UD-PrOZA team at Ghent University Hospital and included SNP-array based homozygosity mapping, Sanger sequencing and cDNA analysis of *BSCL2*, whole exome sequencing (WES), low-pass shallow whole genome sequencing (CNV-sequencing) and whole genome sequencing (WGS). In family 2, genetic analysis was initiated by a clinical geneticist at Hôpital de Hautepierre in Strasbourg and included analysis of neuropathy and lipodystrophy gene panels, WES and Sanger sequencing (Supplementary Methods).

### Transcriptomics

RNA sequencing was performed on an upper arm WAT sample of patient 1 and three healthy lean controls (females, between 30 and 58 years of age and of European Caucasian descent). Online tools Metascape and LISA (epigenetic Landscape In Silico deletion Analysis) were used to perform enrichment analysis of differentially expressed genes (DEG) (Supplementary Methods).

### Lipidomics

Lipidomics analysis was performed on subcutaneous upper inner arm WAT specimens from two patients and 6 control individuals (all female, between 27 and 57 years of age and of European Caucasian descent) in collaboration with Lipometrix™, the lipidomics core at KU Leuven in Belgium. The subcutaneous adipose tissue specimens were obtained through an open biopsy and the site of sampling was identical for all patients and controls. After lipid extraction and sample normalization hydrophilic interaction liquid chromatography-mass spectrometry (HILIC LC-MS/MS) was done, enabling quantification of 1800 lipid species across 16 different lipid classes (https://www.lipometrix.be) (Supplementary Methods).

## Results

### Patients

#### Family 1

Three sisters (patients 1-3), born from consanguineous parents of Turkish origin, presented with lipoatrophy of limbs and trunk, muscle hypertrophy, insulin-resistant diabetes, hypertriglyceridemia with low HDL and liver steatosis (Figure 1A, 2A,C and Table 1). There was an accumulation of adipose tissue in the submental and posterior cervical region in patients 2 and 3 (data not shown). The three patients were lean with a body-mass index (BMI) ranging from 19.6 to 20.5 kg/m^2^. Body fat percentage in patient 1 was 13.8% (Table 1). Serum leptin levels in patient 1 and patient 2 were decreased (Table 1). Additional features included hirsutism, severe acne, androgenic alopecia, polycystic ovary syndrome, acanthosis nigricans, arterial hypertension, subclinical hypothyroidism, precocious puberty and acromegalic features (Figure 2D,E, Table 1 and Supplementary Subjects). Neurological symptoms were also present and comprised severe generalized musculoskeletal pain, migraine, demyelinating peripheral neuropathy and intellectual disability (Table 1). Light microscopic analysis of upper arm subcutaneous WAT in patient 1 showed significantly larger (p=0.016) and more irregularly shaped (p<0.0001) adipocytes with increased inflammation compared to control WAT (Figure 2F-K). There was no difference in the degree of adipose tissue fibrosis (Figure S2).

**Figure 1.**
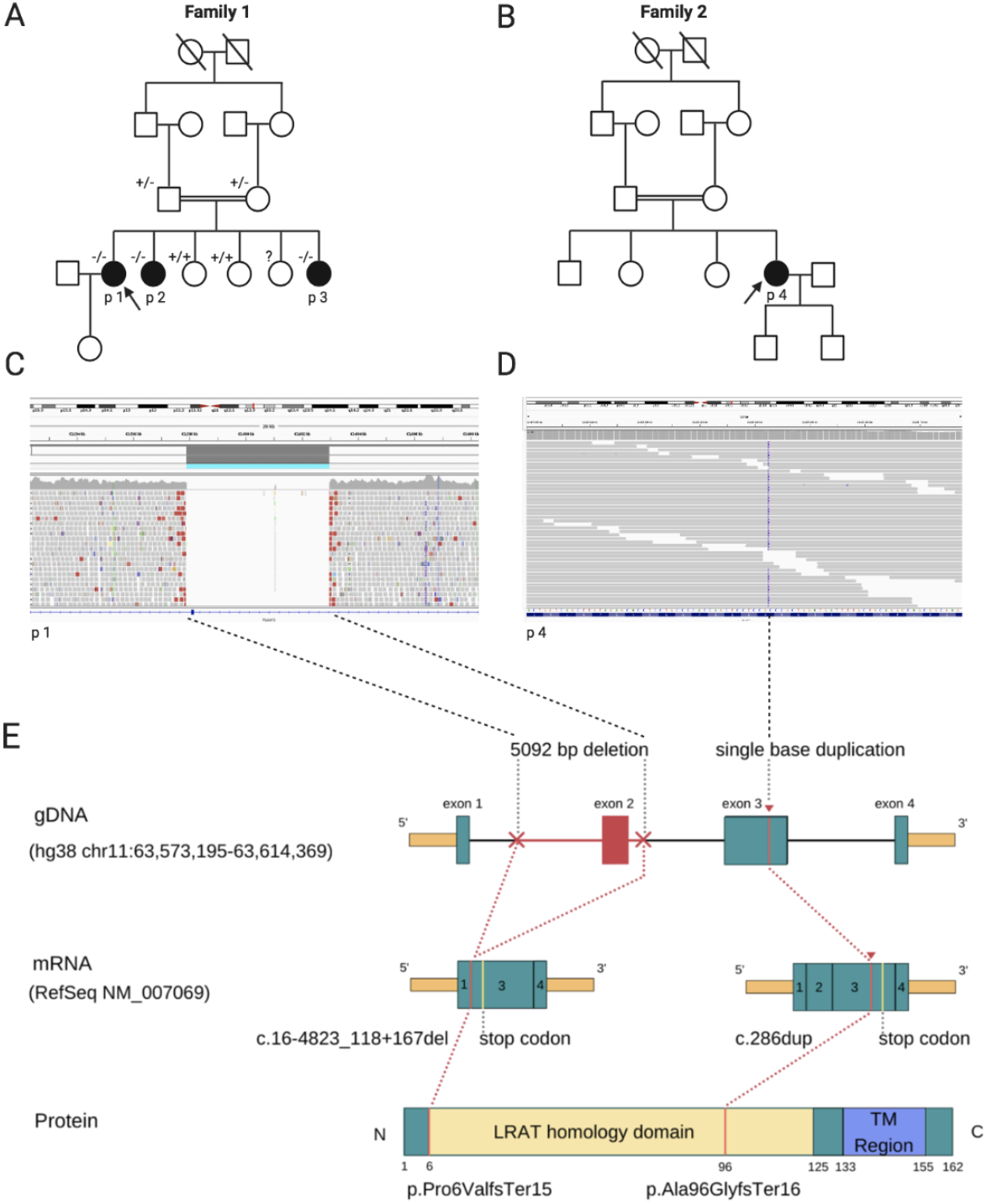
Family pedigree structures, IGV visualization of WGS/WES data and schematic display of the *PLAAT3* LoF mutations in families 1 and 2. **A** Pedigree of family 1. Three affected siblings are born from consanguineous parents who are first cousins of each other. **B** Pedigree of family 2. The proband (patient 4) is the fourth child of consanguineous parents with a first cousin relationship. **C** IGV visualization of WGS reads of patient 1 demonstrates a homozygous deletion of 5092 bp including exon 2 of *PLAAT3*. **D** IGV visualization of WES reads in patient 4 shows a homozygous single base insertion in exon 3 of *PLAAT3*. **E** The homozygous deletion of exon 2 of the *PLAAT3* gene in family 1 and homozygous single base insertion (c.286dup) in exon 3 in family 2 and the consequences of the mutations at the mRNA level are shown. The enzymatically active LRAT and a transmembrane region (TM) of PLAAT3 are depicted. Both mutations result in a frameshift, causing a premature termination codon and are predicted to cause NMD.

**Figure 2.**
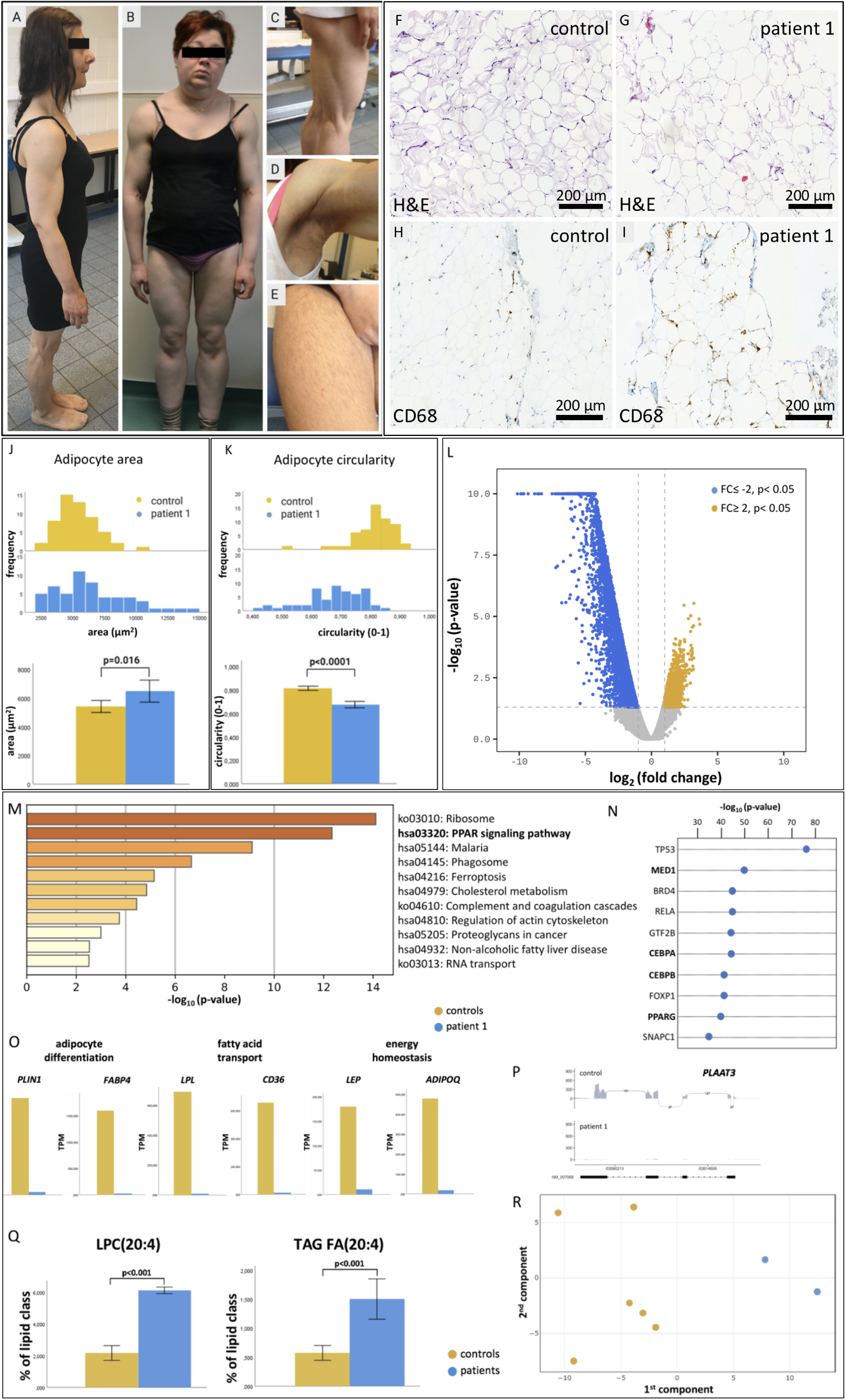
Clinical pictures of patient 1,2,3 and 4, histopathology, transcriptomics and lipidomics analysis of patient and control WAT biopsies. **A** Lateral view of patient 1 showing muscular hypertrophy in upper and lower limbs. **B** Frontal view of patient 4 demonstrating masculine features with muscle hypertrophy and lipoatrophy of upper and lower limbs with a submental accumulation of adipose tissue. **C** Lipoatrophy and muscle hypertrophy of the lower limbs in patient 2. **D** Acanthosis nigricans in the axillary region of patient 2. **E** Hirsutism on the upper lower limb in patient 3. **F**,**G** H&E staining shows the presence of larger and more irregularly shaped adipocytes in patient WAT compared to the control specimen. **H**,**I** CD68 immunohistochemistry showing increased inflammation in patient WAT compared to a control. **J**,**K** Distribution of size (µm^2^) and shape (circularity ranging from 0-1) of the measured adipocytes depicted in histograms. The means are visualized as bar charts with 95% CI error bars. An unpaired T-test (α=0.05) showed significant differences between patient and control adipocytes (p-values are shown above the bars). **L** Volcano plot showing the down-(blue) and upregulated (yellow) genes in patient WAT compared to three controls. **M** KEGG pathway enrichment analysis of top 100 downregulated genes using Metascape. **N** Analysis of transcriptional regulator enrichment of top 500 downregulated genes using LISA. Transcriptional regulators involved in adipocyte differentiation and function are depicted in bold. **O** Mean TPM values of six genes (*PLIN1, FABP4, LPL, CD36, LEP, ADIPOQ*) under transcriptional control of PPARG in controls and patient 1 visualized in bar charts. **P** Sashimi plot visualizing the absence of PLAAT3 transcript in patient WAT. **Q** LPC(20:4) and TAG FA(20:4) content (% of lipid class) in two patient and 6 control samples. The means ± 2 SE are visualized in bar charts. An unpaired student T-test (α=0.05) showed significant differences (p-values are shown above the bars). **R** Principal component analysis (PCA) of lipidomics data shows that patient and control samples cluster separately along the first component.

**Table 1.** Overview of phenotypic features of patients 1,2,3 and 4.

|  | Family 1 |  |  | Family 2 |
| --- | --- | --- | --- | --- |
|  | Patient 1 | Patient 2 | Patient 3 | Patient 4 |
|  | p.Pro6ValfsTer15<br>homozygous | p.Pro6ValfsTer15<br>homozygous | p.Pro6ValfsTer15<br>homozygous | p.Ala96GlyfsTer16<br>homozygous |
| Age (yrs) | 38 | 37 | 25 | 42 |
| Sex | Female | Female | Female | Female |
| Height (cm) | 150 | 153 | 151 | 165 |
| Weight (kg) | 45 | 48 | 51 | 67 |
| Body-mass index (kg/m<sup>2</sup>)<br>(ref: 18.5-25) | 19.7 | 20.5 | 20.5 | 24.6 |
| Fat percentage (%)<br>(ref: 18-30) | 13.8 | NA | NA | NA |
| Triglycerides (mg/dL)<br>(ref: 41-194) | 3745 | 973* | 386 | 205 |
| HDL cholesterol (mg/dL)<br>(ref: 39-96) | 26 | 31* | 31 | 30 |
| Fasting glucose (mg/dL)<br>(ref: 72-110) | 234 | 179* | 129 | 141* |
| HbA1c (%)<br>(ref: 4.0-5.5) | 9.7 | 7.8* | 7.3 | 6.5* |
| Insulin (pmol/L)<br>(ref: 0-174) | 236* | NA | 263 | 232* |
| AST<br>(ref: 0-31 U/L) | 20 | 33 | 16 | 27 |
| ALT<br>(ref: 7-31 U/L) | 17 | 55 | 20 | 38 |
| Gamma-GT<br>(ref: <36 U/L) | 18 | 43 | 20 | 26 |
| CK<br>(ref: <170 U/L) | 52 | NA | 75 | 407 |
| TSH<br>(ref: 0.27-4.20)) | 1.4 | NA | 7.04 | 0.74 |
| Leptin (µg/L)<br>(ref: 3.8-77.6) | <1.6 | 3.5 | NA | 7.4 |
| ***Classical lipodystrophy features*** |  |  |  |  |
| Lipoatrophy of limbs | + | + | + | + |
| Cervico-facial<br>lipohypertrophy | - | + | +/- | + |
| Muscle hypertrophy | + | + | + | + |
| Diabetes mellitus type 2 | + | + | + | + |
| Acanthosis nigricans | - | + | - | + |
| Hirsutism | + | + | + | + |
| Polycystic ovary syndrome | + | NA | + | + |
| Precocious puberty | - | NA | + | - |
| Dyslipidemia | + | + | + | + |
| Hepatomegaly/hepatic steatosis | + | + | + | + |
| ***Additional features*** |  |  |  |  |
| Hypothyroidism/goiter | - | NA | + | + |
| Glomerulopathy | NA | NA | NA | + |
| Hypertension | + | NA | + | + |
| Peripheral demyelinating neuropathy | + | NA | NA | + |
| Intellectual disability | - | + | - | - |
| Chronic muscle/joint pain | + | NA | + | + |

NA: not available, \* data after initiation of treatment

#### Family 2

Patient 4 was born from Algerian first-degree cousins and was the only affected patient in the family (Figure 1B). She presented with an android habitus with generalized muscle hypertrophy and lipoatrophy in the limbs with a relative over-distribution of adipose tissue in the face, neck and submental region resulting in a cushingoid appearance. She complained of chronic muscle pain and was diagnosed with a demyelinating neuropathy (Figure 2B, Table 1). She had a BMI of 24.6 kg/m^2^. Additionally, she had non-insulin-dependent diabetes mellitus, acanthosis nigricans, hypertriglyceridemia with low HDL cholesterol, arterial hypertension, transient hypothyroidism, polycystic ovary syndrome, hirsutism, bilateral carpal tunnel syndrome, obstructive sleep apnea syndrome and a glomerulopathy without renal insufficiency. Serum leptin levels were within the lower normal range (Table 1).

### Genetic studies

#### Family 1

We performed SNP array-based homozygosity mapping in the three affected and two unaffected siblings. The largest homozygous region (∼43 Mb), exclusively shared between the affected siblings, was found on chromosome 11p11.2-q22.1 and contained the *BSCL2* gene (11q12.3), which is associated with congenital generalized lipodystrophy type 2 (CGL2) (Table S1). Normal Sanger sequencing of *BSCL2* coding exons, followed by a normal cDNA analysis on peripheral white blood cells excluded *BSCL2* as the causal gene. WES did not reveal shared pathogenic single nucleotide or small insertion-deletion variants within the shared homozygous regions (Table S2). Using the ExomeDepth algorithm^10^, a homozygous 103 bp deletion corresponding to exon 2 of the *PLAAT3* gene (NM_007069.3) was identified in the three affected sisters but not in the unaffected siblings in whom the deletion was not found (Table S3). The deletion is located within the ∼43 Mb shared homozygous region and was confirmed by low-pass whole genome sequencing (CNV-seq).^11^ Whole-genome sequencing (WGS) in one unaffected and two affected sisters determined the exact genomic deletion breakpoints (chr11: 63597894-63602986) revealing a homozygous 5092 bp deletion (Figure 1C). The absence of WES- and WGS-reads mapped to this genomic region can be visualized using the Integrative Genome Viewer (IGV).^12^ Deletion of *PLAAT3* exon 2 (c.16-4823_118+167del) results in a frameshift leading to a premature termination codon (p.Pro6ValfsTer15). RNA sequencing confirmed that the mutation induces nonsense-mediated decay (NMD) (Figure 1E, Figure 2P).

#### Family 2

Using the GeneMatcher exchange platform^13^, we identified a female patient (patient 4) in which WES identified a homozygous single base duplication in exon 3 of *PLAAT3* leading to a frameshift and premature termination codon, predicted to induce NMD (c.286dupG, p.Ala96GlyfsTer16) (Figure 1D,E). The mutation was confirmed with Sanger sequencing (Figure S1).

Neither of the loss-of-function mutations identified in these two families was present in the gnomAD v2.1.1 database, consisting of 125,748 exomes and 15,708 genomes, nor did it contain other *PLAAT3* loss-of-function mutations in the homozygous state. WGS analysis of 26 additional unexplained lipodystrophy patients identified a causal defect in known lipodystrophy genes in six patients, no additional *PLAAT3* cases were identified (Table S4).

### Transcriptomics

We performed RNA sequencing in WAT samples of patient 1 and three healthy controls in order to identify DEGs as a consequence of PLAAT3 deficiency. Multidimensional scaling showed a marked difference between patient and control transcriptomes (data not shown). A total of 8711 genes were identified which were significantly up- or downregulated in patient WAT (|fc| ≥ 2, raw p<0.05) (data not shown). Of these 8711 genes, the large majority (n=6710) was downregulated in patient WAT (Figure 2L). KEGG pathway enrichment analysis of the 100 most strongly downregulated genes (protein coding and non-coding) identified “ribosome” and “PPAR signaling pathway” as the most enriched clusters (Figure 2M). Next, we used the LISA tool to identify transcriptional regulators (TRs) of gene networks within in the 500 most strongly downregulated genes. Interestingly, out of the top 10 ranked TRs by LISA, four were strongly linked to PPAR signaling including PPARG, the master regulator of adipocyte biology, its two main cooperating TRs CEBPA and CEBPB and its co-activator MED1 (Figure 2N).^14, 15^ In line with these results, transcript levels of well-established PPARG target genes involved in adipocyte differentiation (*PLIN1, FABP4)*, fatty acid transport *(LPL, CD36)* and energy homeostasis (*LEP, ADIPOQ*) were almost completely suppressed in patient adipocytes compared to the controls (Figure 2O). These results indicate a PPARG-mediated adipogenesis defect in PLAAT3-deficient WAT.

### Lipidomics

To study the biochemical consequences of PLAAT3 deficiency in human WAT, we performed LC-MS/MS based lipidomics analysis on a WAT sample of two patients (patient 1 and 4) and compared it to 6 controls. Principal component analysis (PCA) based on all the measured lipid species across all classes showed that the lipid profile in the patients clearly differed from the controls (Figure 2R). As expected, triacylglycerol (TAG) was the most abundant lipid class in patient as well as control WAT samples (Figure S3). The relative abundance of the 16 measured lipid classes was comparable in patient and control WAT (Figure S4). Within the lipid classes however significant differences were observed (Table S5). PLAAT3-deficient WAT showed a higher content of lysophosphatidylcholines (LPC) enriched for arachidonic acid (AA, 20:4) (Log2(FC) =1.5±0.16, p<0.001) (Figure 2Q). Within the TAG class, there were several species for which a statistically significant difference was seen between the patient and control samples (Table S5) including the AA-containing species (Log2(FC)= 1.39±0.34, p<0.001) (Figure 2Q). These results point to an abnormal retention of AA in both the LPC and TAG fractions of PLAAT3 deficient WAT.

## Discussion

The present study links genetic deficiency of phospholipid modifier PLAAT3 to lipodystrophy in humans. Genetic lipodystrophies are rare diseases characterized by reduced WAT, insulin resistance, dyslipidemia and a reduced life expectancy due to an increased cardiovascular risk.^16^ Their general biomedical interest relates to the fact that they are crucial to understand the molecular mechanisms underlying obesity and the metabolic syndrome.^17^ Lipodystrophy patients show striking metabolic similarities with obese subjects due to the incapacity to safely store surplus energy in overwhelmed WAT resulting in ectopic lipid accumulation. Genetic lipodystrophies have been linked to nine different genes and are classified as either congenital generalized lipodystrophy (CGL) [MIM: PS608594], or as familial partial lipodystrophy (FPLD) [MIM: PS151660], depending on the severity and the distribution of the lipoatrophy. PLAAT3 deficiency in our patients was associated with pronounced lipodystrophy of the limbs, with a variable excess of adipose tissue in the facial and cervical regions (patient 2, 3 and 4). Since the phenotypes in our families are most consistent with FPLD, we establish bi-allelic LoF mutations in *PLAAT3* as a novel cause of autosomal recessive FPLD.

Interestingly, our transcriptomics data in patient-derived PLAAT3-deficient WAT revealed an inactivated PPARG-mediated gene network. PPARG is the master regulator of adipocyte development and function and was identified as the causal gene for autosomal dominant FPLD3.^18^ It was shown that the level of residual PPARG activity correlates with the severity of the lipodystrophy phenotype with homozygotes for pathogenic mutations displaying a severe CGL phenotype.^19^ FPLD3 patients present with post puberty partial lipodystrophy, diabetes, hypertension, hypertriglyceridemia and features of hyperandrogenism including hirsutism and polycystic ovaries. Interestingly, there is a striking female preponderance in FPLD3 indicating that sex hormones might influence PPARG activity. The patients in this study were all female and show strong phenotypic similarities with FPLD3 patients. It is currently unclear however if the neurological features (demyelinating neuropathy and chronic pain) seen in patients 1 and 4 are secondary to the diabetes or a primary part of the phenotype which cannot be excluded since PLAAT3 is also expressed in human brain and peripheral nerves (https://www.gtexportal.org/home/gene/PLA2G16).

To date, a high-affinity endogenous and physiologically relevant ligand for PPARG has not yet been identified but it was shown that naturally occurring compounds including polyunsaturated fatty acids such as AA and AA-derived eicosanoids efficiently modify PPARG activity.^20^ An LC-MS-based metabolomics approach identified AA as a candidate natural PPARG ligand in brain, liver and adipose tissue.^21^ It is therefore of interest that the lipidomic profile in PLAAT3-deficient WAT showed increased levels of AA in the phospholipid LPC fraction in line with its known PLA2 activity to liberate AA from the *sn2* position of PC. Somewhat unexpectedly, AA was also increased in the lipid droplet TAG fraction where PLAAT3 has no known hydrolytic activity. This observation can be explained by recent insights suggesting that lipid droplet TAGs can act as a buffer to ensure lipid homeostasis by avoiding excess toxic accumulation of LPCs within membranes.^22^ As such, PLAAT3 appears to play a role as a PPARG ligand generator by liberating AA from membrane phosphatidylcholines in WAT, thereby stimulating transcription of PPARG-regulated genes necessary for adipogenesis, fatty acid transport and glucose homeostasis. Since our patients are predicted to be deficient of endogenous PPARG ligand, in theory they could benefit from treatment with thiazolidinediones (TZDs), a class of anti-hyperglycemic drugs including rosiglitazone, pioglitazone and troglitazone.

Interestingly, *Plaat3-*null mice have been reported to be resistant to diet-induced obesity and lipodystrophic due to chronically upregulated lipolysis.^4^ The inhibitory effect of Plaat3 on lipolysis was suggested to result from the production of PGE2 through the release of AA from the *sn*-2 position of phospholipids by its PLA2 activity.^2, 4^ As a result, *Plaat3-*null mice had decreased PGE2 levels, abolishing the prostaglandin EP3 receptor-mediated negative feedback on the enzyme adenylyl cyclase leading to increased cAMP levels and continuously stimulated lipolysis.^4^ In contrast, our data in human patients did not reveal evidence for increased lipolysis but indicated a primary PPARG-mediated defect in adipogenesis.

Finally, PLAAT3 was identified as an important picornavirus host factor that is required for viral genome delivery to the cytoplasm.^5, 23, 24^ PLAAT3 deficient mice demonstrated resistance to picornavirus infection.^5^ Whether lipodystrophy patients with genetic PLAAT3 deficiency are protected against picornavirus infections, needs to be further investigated. Due to its potential anti-obesity and anti-viral properties, modifying PLAAT3 activity might be an interesting therapeutic strategy. Interestingly, while it was shown that drug targets with human genetic support are twice as likely to lead to approved drugs^25, 26^, our study suggests that blocking PLAAT3 activity too strongly could lead to unwanted metabolic side-effects secondary to a downregulation of PPARG-regulated gene networks.

In conclusion, we report four patients from two families harboring bi-allelic loss-of-functions mutations in *PLAAT3*, presenting with partial lipoatrophy, diabetes and hypertriglyceridemia, thereby establishing PLAAT3 deficiency as a novel cause of FPLD. By using a multi-omics strategy, we identified AA-dependent decreased PPARG activity as the underlying mechanism causing adipocyte dysfunction and consequently disturbed energy homeostasis and as such pinpoint PPARγ agonist TZD as a potential therapeutic strategy in these patients.

## Supplemental data

Supplemental data include additional data on subjects, methods, 3 supplemental figures and 5 supplemental tables.

## Acknowledgements

The authors would like to thank the patients and families who participated in this study, the GenomEast facility (IGBMC, Strasbourg, France) for exome sequencing in patient 4. We thank Machteld Baetens for in-depth CNVseq analysis. B.D. is supported by an Odysseus type 1 Grant of the Research Foundation Flanders (3G0H8318) and a starting grant from Ghent University Special Research Fund (01N10319). The Program for Undiagnosed Diseases (UD-PrOZA) is supported by the Spearhead Research Policy Program and the Fund for Innovation from the Ghent University Hospital. The Neuromendeliome Study (to C.D., Strasbourg, France) was financially supported by Agence de la Biomédecine (France).

## Web resources

GeneMatcher, https://www.genematcher.org/

gnomAD, http://gnomad.broadinstitute.org/

Genbank, https://ncbi.nlm.nih.gov/genbank/

OMIM, http://www.omim.org/

Genotype-Tissue Expression (GTEx) project, https://gtexportal.org/

