## Supplemental Data for "Loss of adipocyte phospholipase gene *PLAAT3* causes lipodystrophy and insulin resistance due to inactivated arachidonic acid-mediated PPAR*γ* signaling"

1. **Subjects and methods**
2. **Supplemental figures**
3. **Supplemental tables**

1. **Subjects and methods**

*Subjects*

Family 1 was referred to the Center for Medical Genetics Ghent (Ghent University Hospital) by an endocrinologist for genetic counseling and diagnostic genetic testing. After negative routine genetic testing the family was included in the multidisciplinary Program for Undiagnosed Diseases (UD-PrOZA) at the Ghent University Hospital. The main objective of UD-PrOZA is to solve rare undiagnosed disorders by combining broad clinical expertise (internal medicine, neurology, pediatrics, clinical genetics) with state-of-the art genomic approaches.

- Patient 1 (°1981) is the proband of this family. She was born from parents who are first-degree cousins of Turkish origin. The family history was negative aside from diabetes mellitus type 2 in the mother. She attained menarche at the age of 12 years. At the age of 19, she consulted a physiotherapist because of chronic musculoskeletal pain in upper and lower limbs and a progressive muscular hypertrophy. Around the same time, she started to experience episodes of disabling headaches. At 25 years of age, she consulted an endocrinologist because of a persistent low BMI of 17.3 kg/$m^2$ (39 kg, 150 cm). Diagnostic tests to exclude malabsorption syndromes were negative. In her late twenties she had a spontaneous abortion at a gestational age of 8 weeks. Due to subfertility (for which no clear cause could be identified), IVF was performed after which she gave birth to a healthy child at the age of 30. In the course of the pregnancy, she developed gestational diabetes. At the age of 32, she was diagnosed with a carpal tunnel syndrome at the right side for which a surgical release was performed. At the age of 34, she was diagnosed with diabetes mellitus after a routine blood test revealed a severe hypertriglyceridemia (3745 mg/dL, reference: 41-194 mg/dL) with low HDL (26 mg/dL, reference: 39-96 mg/dL) and hyperglycemia (fasting glucose 234 mg/dL, reference: 72-110 mg/dL and HbA1c 9,7%, reference: 4.0-5.5%). She had very low leptin levels (<1.6 µg/L, reference: 3.8-77.6 µg/L). Microalbuminuria was present but was corrected upon normalization of blood glucose levels. The creatinine clearance and thyroid function were normal. The liver transaminase levels were transiently elevated and 25-OH-Vitamin D levels were consistently low. Abdominal ultrasound and liver elastography showed hepatomegaly with steatosis, saverymuttu score grade 1. At that time, she weighed 45 kg for a body length of 150 cm (BMI of 19,7 kg/$m^2$). A dual X-ray absorptiometry showed a body fat percentage of 13.8%, which is significantly lower than the reference values for women of her age (18-30%). The lipoatrophy was generalized but was less pronounced in the subcutaneous adipose tissue in the upper arms. At the age of 36 years, she started to notice progressive hair loss which was diagnosed as androgenic alopecia. Hirsutism has been present since a young age. The affected sisters can be distinguished from their non-affected sibs by the presence of acromegalic facial features, predominantly reflected in enlarged noses. During the last 10 years the most disabling morbidities from a patient perspective have been an increasing fatigue with exercise intolerance, migraine and upper and lower extremity musculoskeletal pain. MRI of the brain and skeletal muscles were normal. In the medical files, the pain is described to be partly myogenic, partly neurogenic. Electromyoneurography showed the presence of a demyelinating peripheral neuropathy. The headaches are unilateral and are accompanied by photophobia and nausea and are therefore considered to be migrainous, though there was no aura.  The intensity of these headaches reaches the extent that they required multiple emergency department visits for status migrainosus. Recently, she had a cardiac evaluation because of recurrent chest pain. A normal exercise ECG and echocardiography point in the direction of a musculoskeletal origin of the chest pain. She was diagnosed however with an arterial hypertension. Her chronic medication consists of a combinational product of metformin and a DPP4-inhibitor (sitagliptine), Janumet (50mg/1g) two times a day. Additionally, she is treated with Coversyl 5mg/d (perindopril) for hypertension. Cholesterol lowering statin treatment was discontinued because it aggravated her muscle pains.  Instead, she is treated with Lipanthylnano 145 mg/d (fenofibrate). She has worked as a factory worker and a hairdresser in the past, which is currently not possible due to the disabling pain complaints.

- Patient 2 (°1982) is the eldest sister of the proband. She has moderate intellectual disability requiring us to rely on heteroanamnesis. Phenotypically she presents with similar features observed in her sister predominantly consisting of muscle hypertrophy, lipoatrophy, hirsutism and coarse facial features. In the submental region, an excess of adipose tissue is present. In the axillary regions we observed acanthosis nigricans, a skin condition associated with hyperinsulinemia. She has a body-mass index of 20.5 kg/m$^2$. Similar to her sister, she presented with a hypertriglyceridemia (973 mg/dL) with low HDL cholesterol (31 mg/dL) and insulin resistant diabetes mellitus type 2 with high fasting blood glucose levels (179 mg/dL) and elevated HbA1c (7.8%). She had decreased leptin levels (3.5 µg/L).
- Patient 3 (°1994) is the youngest sibling of patient 1 and 2. At 8 years of age, she presented with a central precocious puberty for which she was treated with gonadorelin to maximize her adult height. When she was 18 years old, a routine blood test showed hypertriglyceridemia (386 mg/dL) with a low HDL (29 mg/dL). At the age of 20 years, she started to complain of myalgia in the upper and lower limbs. A progressive lipoatrophy and muscular hypertrophy were noticed. She consulted a dermatologist because of severe facial hirsutism for which laser treatment was performed. Additionally, she had severe acne during adolescence. A blood test at the age of 22 years showed mildly elevated liver enzyme plasma concentrations (AST: 46 U/L, reference: 0-31 U/L; ALT: 73 U/L, reference: 7-31 U/L; gamma-GT: 34 U/L, reference: <36 U/L). She had low 25-OH-Vitamin D levels. She was diagnosed with diabetes mellitus and insulin resistance based on fasting glucose level of 121 mg/dL, HbA1c of 7,3% and hyperinsulinemia of 236 pmol/L (reference: 0-174 pmol/L). At the age of 25, she developed subclinical hypothyroidism (TSH: 7,4 mU/L, reference: 0.27-4.20 mU/L with normal T3 and T4 values) with subtle enlargement of the thyroid gland. She weighs 50,6 kg for a height of 151 cm (BMI: 20,5 kg/m$^2$). She complains of chronic pain in both arms and legs for which no further

diagnostic tests have been performed. There is a generalized absence of adipose tissue, with the exception of the submental and dorsal cervical region where an accumulation of adipose tissue is present. Currently, she is not taking antidiabetic medication. Lifestyle measures suffice to control glycemia levels and Hb1Ac. She works for the Belgian post sorting center, which is manageable despite the chronic limb pain.

Family 2 includes a 42-year-old female patient born from healthy first-degree cousins born in Algeria. She has two healthy sons and has no familial history. She has two sisters living in Algeria who, according to the patient, are asymptomatic. She has an older brother who was seen at the genetics department lacking any clinical signs of lipodystrophy, though genetic analysis is still ongoing. Patient 4 presented at the first evaluation with limb lipoatrophy and muscular hypertrophy, facio-cervical lipohypertrophy, hyperandrogenism including hirsutism in a context of polycystic ovary syndrome, non-insulin-dependent diabetes with acanthosis nigricans, dyslipidaemia (hypertriglyceridemia and low HDL cholesterol), arterial hypertension, thyroid goitre with benign macronodule and demyelinating polyneuropathy. The lipodystrophy appeared during adolescence. The weight was 67 kg for a height of 165 cm (BMI of 24.6 kg/m$^2$). The evolution was marked by a progressive increase of the muscular hypertrophy and of the submental lipohypertrophy with several episodes of respiratory discomfort. Chronic and generalized muscular pain and weakness appeared and led progressively to social withdrawal. She was diagnosed with recurrent carpal tunnel syndrome for which several infiltrations with steroids were given. She developed a sleep apnea syndrome and polysomnography showed severe obstructive apnoea-hypopnea requiring nocturnal ventilation. In addition, a glomerulopathy was diagnosed, probably due to focal segmental glomerulosclerosis (no renal biopsy performed), without renal failure. Cardiovascular assessment showed isolated arterial hypertension. Laboratory tests showed hypertriglyceridemia (205 mg/dL), low HDL cholesterol (30 mg/dL), normal liver enzymes and transient hypothyroidism. She had elevated insulin levels (232 pmol/L) and leptin levels close to the lower limit (7.4 µg/L). These analyses were not performed in a fasting condition. Brain MRI was normal. Bone x-rays showed vertebral posterior osteophytes. CK-levels were elevated on one analysis

(470 U/L, reference: <170 U/L), most likely secondary to statin treatment. A muscular biopsy showed mild myogenic anomalies, but no signs of a real myopathy. Cervical CT scan showed an increased amount of fat in the submental region. Lipodystrophy and polyneuropathy gene panels (including *AKT2, BSCL1, BSCL2, CAV1, CIDEC, DYRK1B, INSR, LIPE, LMF1, LMNA, LMNB2, NSMCE2, PYK3R1, PLD3, PLIN1, POC1A, POLD1, PPARG, PSMB8, PTRF, PCYT1A, TBC1D4,* and *ZMPSTE24)* were negative. Currently, she is being treated with semaglutide 0.5 mg (subcutaneous injection 1x/w), metformin 700 mg 2x/d, valsartan 80 mg 1x/d and amlodipine 10 mg 1x/d.

*Genetic studies*

Family 1:

Homozygosity mapping: Genotyping for homozygosity mapping was performed in three affected and two unaffected members of family 1 using 200K genome-wide HumanCytoSNP-12 v2 BeadChip single nucleotide polymorphism (SNP) arrays (Illumina, San Diego, CA). The position of the probes was based on NCBI build GRCh37. Homozygous regions shared between affected members were detected using the PLINK algorithm (v1.07, default settings).[1]

*BSCL2* mutation screening: Sequencing of *BSCL2* was done using Sanger sequencing after PCR amplification of all exons including splice junctions. Extracted mRNA from cultured lymphocytes was retrotranscribed to cDNA. The region covering *BSCL2* exons 2 to 9 were amplified by PCR from this cDNA. The PCR product was migrated on an agarose gel and its size was compared to that obtained of a control subject.

Whole-exome sequencing (WES): WES was done on the Illumina Novaseq 6000 Platform after enrichment of gDNA with SureSelectXT Low Input Human All Exon v7 (Agilent Technologies). The BWA-MEM 0.7.17 algorithm was used for read mapping against the human genome reference sequence (NCBI, GRCh37.p5/hg19), duplicate read removal, and variant calling. Variant calling and filtering were done using Seqplorer, an in-house developed tool for the analysis of WES-data. The position of the called variants is based on NCBI build GRCh38. A minimum of 90% of the interrogated genes have a coverage of >20x. Variant classification was done according to the ACMG guidelines.[2] CNVs were detected with ExomeDepth.[3]

Molecular karyotyping was done by means of low-pass whole-genome sequencing (CNV-seq) on the Illumina Novaseq 6000 with a genome wide resolution of 100kb (GRCh38).

Whole-genome sequencing was performed by Macrogen on an Illumina Novaseq 6000 platform. Fastq-files were aligned against the hg38 reference genome with BWA-MEM (v0.7.17). Files were converted to BAM-format, duplicate marked, and sorted and indexed using Samtools (v1.9)[4] and Picard (v2.21.6). Structural variants (SVs) were then called using three different callers, namely DELLY (v0.8.3)[5], LUMPY (v0.2.13)[6] and Manta (v1.6.0)[7]. Standard settings were used for all.

RNA paired-ends sequencing was performed by Macrogen on an Illumina platform. Library preparation was done using the SMARTer Universal Low Input RNA Kit and the TruSeq RNA Sample Prep Kit v2. Trimmed reads were mapped to the reference genome with HISAT2.[8] After read mapping, Stringtie was used for transcript assembly.[9] Expression profile was calculated for each sample and transcript/gene as read count, FPKM (Fragment per Kilobase of transcript per Million mapped reads) and TPM (Transcripts per Kilobase Million). DEG (Differentially Expressed Genes) analysis was performed on a comparison pair (Test vs Control) using edgeR.[10] Functional enrichment analysis of KEGG pathways[11] within 100 most strongly downregulated genes was performed using the Metascape tool[12]. To identify potential transcriptional regulators (TRs) of the top 500 downregulated genes we performed LISA (epigenetic Landscape In Silico deletion Analysis; http://lisa.cistrome.org/), which is designed to combine a comprehensive database of human and mouse DNase-seq, H3K27ac ChIP-seq, as well as TR Chip-seq to TRs that regulate a query gene set.[13]

Family 2:

WES was done for the index case (patient 4) as part of the 'Neuromendeliome' study, including 26 patients with unsolved syndromes affecting the nervous system. Paired-end sequencing libraries were prepared using the Agilent SureSelect XT Human All Exon v7 Enrichment kit. Sequencing (2x100 bases) was performed on a HiSeq2500 (Illumina) on the GenomEast platform (IGBMC, Illkirch, France). Image analysis and base calling were performed using CASAVA v1.8.2 (Illumina). Reads were mapped onto the reference genome Hg19 using BWA v0.7.5a.[14] Elimination of duplicate reads and base quality recalibration were performed using Picard v1.122. Realignment around indels was done with GATK v3.2-2. Reads mapping to several positions in the genome were excluded using Samtools v0.1.19.[4] Variant calling was done using GATK v3.2-2 Unified genotyper.[15] Variants are annotated using GATK v3.2-2[15], SnpE_ v2.0.5[16] and SnpSift v4.4l.[17]

*Lipidomics*

Lipid extraction

An amount of sample containing 10 µg of protein was mixed with 800 µl 1 N HCl:CH3OH 1:8 (v/v), 900 µl CHCl3, 200 µg/ml of the antioxidant 2,6-di-tert-butyl-4-methylphenol (BHT; Sigma Aldrich) and 3 µl of SPLASH® LIPIDOMIX® Mass Spec Standard (#330707, Avanti Polar Lipids). After vortexing and centrifugation, the lower organic fraction was collected and evaporated using a Savant Speedvac spd111v (Thermo Fisher Scientific) at room temperature and the remaining lipid pellet was stored at -20°C under argon.

Mass spectrometry

Just before mass spectrometry analysis, lipid pellets were reconstituted in 100% ethanol. Lipid species were analyzed by liquid chromatography electrospray ionization tandem mass spectrometry (LC-ESI/MS/MS) on a Nexera X2 UHPLC system (Shimadzu) coupled with hybrid triple quadrupole/linear ion trap mass spectrometer (6500+ QTRAP system; AB SCIEX). Chromatographic separation was performed on a XBridge amide column (150 mm × 4.6 mm, 3.5 µm; Waters) maintained at 35°C using mobile phase A [1 mM ammonium acetate in water-acetonitrile 5:95 (v/v)] and mobile phase B [1 mM ammonium acetate in water-acetonitrile 50:50 (v/v)] in the following gradient: (0-6 min: 0% B → 6% B; 6-10 min: 6% B → 25% B; 10-11 min: 25% B → 98% B; 11-13 min: 98% B → 100% B; 13-19 min: 100% B; 19-24 min: 0% B) at a flow rate of 0.7 mL/min which was increased to 1.5 mL/min from 13 minutes onwards. SM, CE, CER, DCER, HCER, LCER were measured in positive ion mode with a precursor scan of 184.1, 369.4, 264.4, 266.4, 264.4 and 264.4 respectively. TAG, DAG and MAG were measured in positive ion mode with a neutral loss scan for one of the fatty acyl moieties. PC, LPC, PE, LPE, PG, PI and PS were measured in negative ion mode by fatty acyl fragment ions. Lipid quantification was performed by scheduled multiple reactions monitoring (MRM), the transitions being based on the neutral losses or the typical product ions as described above. The instrument parameters were as follows: Curtain Gas = 35 psi; Collision Gas = 8 a.u. (medium); IonSpray Voltage = 5500 V and −4,500 V; Temperature = 550°C; Ion Source Gas 1 = 50 psi; Ion Source Gas 2 = 60 psi; Declustering Potential = 60 V and −80 V; Entrance Potential = 10 V and −10 V; Collision Cell Exit Potential = 15 V and −15 V.

The following fatty acyl moieties were taken into account for the lipidomic analysis: 14:0, 14:1, 16:0, 16:1, 16:2, 18:0, 18:1, 18:2, 18:3, 20:0, 20:1, 20:2, 20:3, 20:4, 20:5, 22:0, 22:1, 22:2, 22:4, 22:5 and 22:6 except for TGs which considered: 16:0, 16:1, 18:0, 18:1, 18:2, 18:3, 20:3, 20:4, 20:5, 22:2, 22:3, 22:4, 22:5, 22:6.

Data Analysis

Peak integration was performed with the MultiQuant™ software version 3.0.3. Lipid species signals were corrected for isotopic contributions (calculated with Python Molmass 2019.1.1) and were quantified based on internal standard signals and adheres to the guidelines of the Lipidomics Standards Initiative (LSI) (level 2 type quantification as defined by the LSI). Unpaired T-test p-values and FDR corrected p-values (using the Benjamini/Hochberg procedure) were calculated in Python StatsModels version 0.10.1.


### 2. Supplemental figures

**Figure S1.** Confirmation of single base insertion in exon 3 of *PLAAT3* gene with Sanger sequencing on cDNA of patient 4.

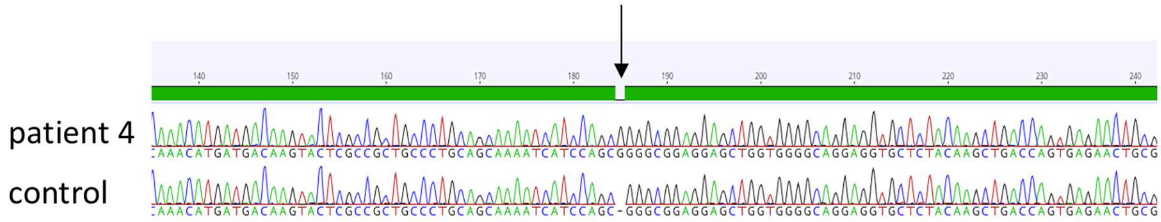


**Legend:**

WES in patient 4 showed the presence of a single base insertion in exon 3 of the PLAAT3 gene (c.286dup). The presence of a homozygous single base insertion was confirmed by Sanger sequencing. The inserted nucleotide is indicated with an arrow.

**Figure S2.** Light microscopic analysis of control and patient WAT after staining with Sirius red.

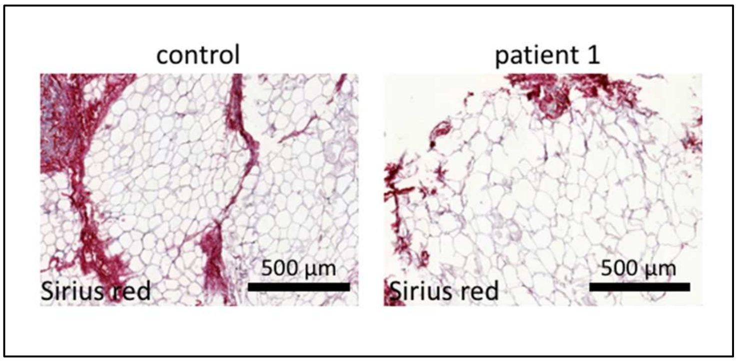


**Legend:**

Sirius red staining and light microscopy of control and patient adipose tissue showing no difference in the degree of fibrosis.

**Figure S3.** Pie chart visualization of relative abundance of all measured lipid classes in a WAT biopsy of patients 1 and 4 compared to 6 controls.

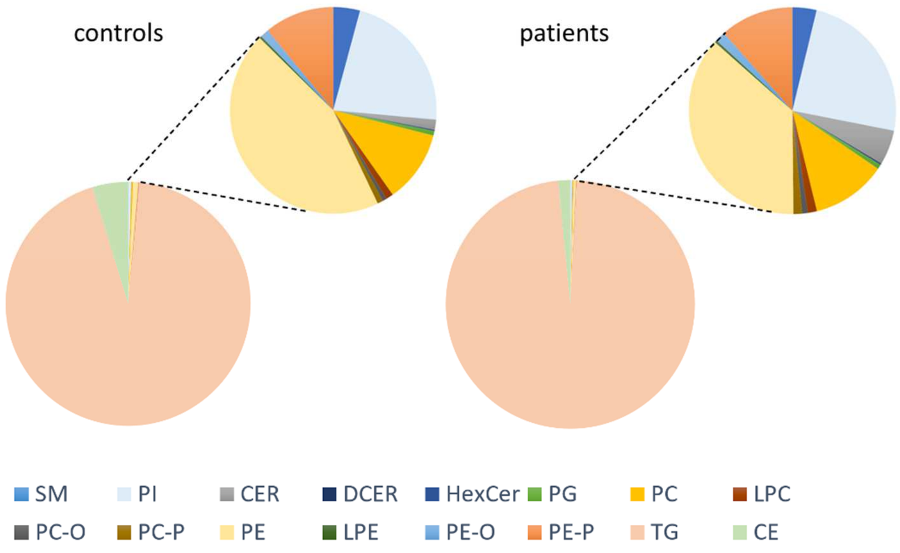


**Legend:**

Distribution of 16 lipid classes, as percentages of total lipid content, visualized as pie charts. The most abundant lipid class, as expected, is the class of the triacylglycerides (TAG) accounting for 94% and 98% of the total lipid content in controls and patients respectively. (SM: sphingomyelins, PI: phosphatidylinositol, CER: ceramides, DCER: dihydroceramides, HexCer: hexosylceramides, PG: phosphatidylglycerol, PC: phosphatidylcholine, LPC: lysophosphatidylcholine, PC-O: 1-alkyl,2-acylphosphatidylcholine, PC-P: 1-alkenyl,2-acylphosphatidylcholine, PE: phosphatidylethanolamine, LPE: lysophosphatidylethanolamine, PE-O: 1-alkyl,2-acylphosphatidylethanolamines, PE-P: 1-alkenyl,2-acylphosphatidylethanolamines, TG: triacylglycerides, CE: cholesterol esters)

### 3. Supplementary Tables

**Table S1**. Homozygosity mapping in family 1.

| Homozygous regions shared between patients 1,2 and 3 (GRChg37) | Cytoband | Size | Nr. probes | Nr. genes | Known CL candidate genes |
|---|---|---|---|---|---|
| Chr1:32,183,736-32,729,702 | 1p35.1-p35.2 | 545,967 | 73 | 17 | - |
| Chr8:91,438,561-92,586,901 | 8q21.3-q22.1 | 1,148,341 | 161 | 10 | - |
| Chr11:44,393,273-87,467,954 | 11p11.2-q14.2 | 43,074,682 | 3669 | 702 | *BSCL2* |
| Chr13:69,045,126-73,179,907 | 13q21.33-q22.1 | 4,134,782 | 412 | 6 | - |
| Chr13:80,259,570-94,062,868 | 13q31.1-q31.3 | 13,803,299 | 900 | 42 | - |

**Table S2.** Rare (MAF < 2%) homozygous variants detected in shared homozygous regions in patients 1,2 and 3 through WES

| Gene | Cytoband | RefSeq Transcript | Variant c-notation | Variant p-notation | Allele frequency* | Class** |
|---|---|---|---|---|---|---|
| ***B3GAT3*** | 11q12.3 | NM_012200.3 | c.979C>T | p.Arg327Trp | 0.03% | Class 3 |
| ***IGHMP2*** | 11q13.3 | NM_002180.2 | c.767C>G | p.Ala256Gly | 0.1% | Class 3 |
| ***TPCN2*** | 11q13.3 | NM_139075.3 | c.699C>T | p.Phe233= | 0.06% | Class 1 |

*European (non-Finnish) population

**Based on ACMG-guidelines

**Table S3**. CNV analysis in family 1 using WES (ExomeDepth) and low-pass WGS (CNV-seq) data showing potential CNVs present in at least 2 of the three affected sisters.

| | | ExomeDepth | | | | | | | CNV-seq |
|---|---|---|---|---|---|---|---|---|---|
| Cytoband | Gene | Genomic region (hg38) | CNV size (bp) | Number of exons in CNV | Patient Nr | Reads expected | Reads Observed (Patient) | Reads Ratio | CNV type | Confirmation (Y/N) |
| 2q34 | *UNC80* | chr2:209849451-209970957 | 121507 | 32 | 2 | 4350 | 2386 | 0.55 | del | Y |
| | | | | | 3 | 5203 | 2959 | 0.57 | del | Y |
| 3p21.2 | *DOCK3* | chr3:50841675-50841715 | 41 | 1 | 1 | 75 | 133 | 1.77 | dup | N |
| | | | | | 3 | 87 | 187 | 2.15 | dup | N |
| 7q22.1 | *SPDYE3* | chr7:100311750-100314750 | 3001 | 3 | 1 | 74 | 17 | 0.23 | del | N |
| | | | | | 3 | 79 | 15 | 0.19 | del | N |
| **11q12.3** | ***PLAAT3*** | **chr11:63598061-63598163** | **103** | **1** | **1** | **70** | **0** | **0.00** | **del** | **Y** |
| | | | | | **2** | **70** | **0** | **0.00** | **del** | **Y** |
| | | | | | **3** | **76** | **0** | **0.00** | **del** | **Y** |
| 14q11.2 | *DHRS4L2* | chr14:24000863-24001084 | 222 | 2 | 2 | 387 | 559 | 1.44 | dup | N |
| | | | | | 3 | 511 | 864 | 1.69 | dup | N |
| 16p13.11 | *NOMO1* | chr16:14838407-14846683 | 8277 | 4 | 2 | 254 | 101 | 0.40 | del | N |
| | | | | | 3 | 316 | 184 | 0.58 | del | N |
| | | chr16:14857217-14857322 | 106 | 1 | 2 | 238 | 129 | 0.54 | del | N |

| | | | | | | | | | | |
|---|---|---|---|---|---|---|---|---|---|---|
| | | | | | 3 | 316 | 184 | 0.58 | del | N |
| | | chr16:14894998-14895090 | 93 | 1 | 2 | 161 | 82 | 0.51 | del | N |
| | | | | | 3 | 193 | 112 | 0.58 | del | N |
| 16p11.2 | *AC136428.1* | chr16:33844784-33845229 | 446 | 2 | 1 | 646 | 414 | 0.64 | del | N |
| | | | | | 2 | 595 | 371 | 0.62 | del | N |

**Table S4.** WGS analysis of 26 unexplained lipodystrophy cases yielded 6 molecular diagnoses.

| ID | Gene | Mutation | Homozygous/ heterozygous | Diagnosis, inheritance (MIM) |
|---|---|---|---|---|
| **1** | *AGPAT2* | Deletion exon 1 | Homozygous | CGL type 1, AR (608594) |
| **2** | *BSCL2* | c.864-1G>A | Homozygous | CGL type 2, AR (269700) |
| **3** | *CAVIN1* | c.135del, p.Lys45AsnfsTer5 | Homozygous | CGL type 4, AR (613327) |
| **4** | *CAVIN1* | c.520G>T, p.Glu174Ter | Homozygous | CGL type 4, AR (613327) |
| **5** | *LMNA* | c.1445G>A, p.Arg782Gln | Heterozygous | FPLD type 2, AD (151660) |
| **6** | *LMNA* | c.406G>C, p.Asp136His | Heterozygous | FPLD type 2, AD (151660) |

**Table S5.** Lipid species with significant differences (FDR adjusted p-value<0.01) between patients and control samples.

| Lipid ID | Lipid class | Log FC | p-value* | FDR adjusted p-value** |
|---|---|---|---|---|
| **TAG(58:8/FA20:4)** | Triacylglycerides | 2.366 | 0.00007 | 0.008 |
| **TAG(56:7/FA20:3)** | Triacylglycerides | 2.129 | 0.00003 | 0.006 |
| **TAG(56:5/FA16:0)** | Triacylglycerides | 2.042 | 0.00004 | 0.006 |
| **TAG(54:6/FA20:4)** | Triacylglycerides | 1.834 | 0.00007 | 0.008 |
| **TAG(52:3/FA20:3)** | Triacylglycerides | 1.635 | 0.00003 | 0.006 |
| **TAG(54:5/FA20:3)** | Triacylglycerides | 1.524 | 0.000003 | 0.001 |
| **TAG(54:6/FA20:3)** | Triacylglycerides | 1.513 | 0.00003 | 0.006 |
| **LPC(20:4)** | Lysophosphatidylcholine | 1.495 | 0.00009 | 0.009 |
| **LPE(16:1)** | Lysophosphatidylethanolamine | -0.986 | 6e-7 | 0.0006 |

*Unpaired T-test **Benjamini-Hochberg procedure
